# Hormetic heat shock activates HLH-30/TFEB independently of canonical nutrient-sensing pathways

**DOI:** 10.64898/2026.09.01.748694

**Authors:** Tatiana M. Moreno, Michelle E. Brown, Caitlin M. Lange, Gavin McLaren, Diego A. Hernandez-Urbina, Cheng-Ju Kuo, Caroline Kumsta

## Abstract

In *Caenorhabditis elegans*, a brief, sublethal heat shock (HS) induces a hormetic response that increases resistance to subsequent stress and extends lifespan. These benefits require *hlh-30*, the ortholog of mammalian transcription factor EB (*TFEB*). Although HS induces robust HLH-30 nuclear translocation, how this response is regulated remains poorly understood. Nutrient- and energy-sensing pathways, including mTORC1 and AMPK, regulate HLH-30/TFEB subcellular localization under other physiological conditions, but whether they mediate its nuclear translocation during HS is unknown. Here, we show that, although HS inhibited mTORC1 and dephosphorylated its conserved HLH-30 S201 target site, HLH-30 S201 phosphorylation was dispensable for HS-induced HLH-30 nuclear localization and hormetic protection. Moreover, HS remained protective in *hlh-30* mutants when mTORC1 activity was reduced, revealing an HLH- 30-independent component of the hormetic response. HS also activated AMPK and *aak-2* was required for hormetic protection but dispensable for HS-induced HLH-30 nuclear localization and autophagosome formation. Together, these findings demonstrate that although HS engages canonical mTORC1 and AMPK signaling, these pathways do not account for HS-induced HLH- 30 nuclear localization and instead make distinct contributions to hormetic protection. Our findings reveal stress-specific regulation of HLH-30/TFEB and point to additional mechanisms that drive its activation during heat stress.

## INTRODUCTION

Exposure to mild stress can induce adaptive responses that increase resistance to subsequent stress and promote longevity, a phenomenon known as hormesis [1, 2]. In *Caenorhabditis elegans*, a brief heat shock (HS) early in adulthood increases thermotolerance and extends lifespan [3–6]. These benefits require the transcription factor *hlh-30*, the *C. elegans* ortholog of mammalian transcription factor EB (*TFEB*), as well as autophagy [5, 6].

HLH-30/TFEB is a conserved transcriptional regulator of autophagy, lysosomal function, metabolism, and stress responses [7, 8]. In mammalian cells, TFEB activity is regulated in large part through its subcellular localization, which is controlled by post-translational modifications, particularly phosphorylation [9, 10]. We previously showed that hormetic HS induces HLH-30 nuclear localization and increases HLH-30-dependent expression of select autophagy genes [6]. However, how HS induces HLH-30 nuclear translocation and how regulation of HLH-30 subcellular localization contributes to the beneficial effects of hormetic HS remain unknown [11]. Defining how different stresses regulate HLH-30/TFEB is important for understanding how cells adapt their autophagic and lysosomal clearance responses and improve cellular homeostasis to different physiological challenges.

One of the major regulators of TFEB localization is mechanistic target of rapamycin complex 1 (mTORC1). Under nutrient-rich conditions, mTORC1 phosphorylates TFEB at several serine residues, including S122, S142, and S211 [9, 12–14]. Among these, phosphorylation of S211 plays a major role in TFEB cytoplasmic retention by promoting binding to 14-3-3 proteins [12, 13]. Upon mTORC1 inhibition, TFEB is dephosphorylated and translocates into the nucleus [9, 12]. This regulation is at least partially conserved in *C. elegans*. Inhibition of *let-363*/*mTOR* by RNAi promotes HLH-30 nuclear localization and autophagy gene transcription [8], and substitution of HLH-30 S201 with alanine, corresponding to human TFEB S211, promotes partial HLH-30 nuclear localization under basal conditions [15]. mTORC1 inhibition, through RNAi against mTORC1 components or rapamycin treatment, also increases stress resistance and lifespan in *C. elegans* [16, 17]. Since these effects overlap with those induced by hormetic HS, mTORC1 inhibition provides a potential mechanism linking HS to HLH-30 activation and hormetic protection.

AMP-activated protein kinase (AMPK) is conserved energy-sensing kinase that could contribute to the regulation of HLH-30 during HS. AMPK is activated in response to low cellular energy and promotes stress adaptation by inhibiting energy-consuming processes and activating pathways that restore cellular homeostasis [18]. In mammalian cells AMPK promotes TFEB nuclear localization and transcriptional activity during energetic stress, although the underlying mechanisms appear to be context dependent [19–21]. AMPK can also induce autophagy through phosphorylation and activation of ULK1/UNC-51, an upstream regulator of autophagosome formation [22, 23]. In *C. elegans*, constitutive activation of AAK-2/AMPK extends lifespan [24], and *aak-2* is required for lifespan extension upon hormetic HS [25]. HS also reduces cellular energy levels, consistent with activation of AAK-2 under these conditions [25]. Thus, AMPK could contribute to hormetic protection through regulation of HLH-30, autophagy, or both.

While mTORC1 and AMPK are well-established regulators of cellular responses to nutrient and energy availability, whether these pathways regulate HLH-30/TFEB during HS is unclear. HS imposes both proteotoxic and energetic stress [25, 26], potentially engaging mechanisms of HLH-30/TFEB regulation distinct from those operating during nutrient stress. We therefore asked whether canonical mTORC1- and AMPK-dependent mechanisms account for HLH-30 nuclear localization and hormetic protection in *C. elegans*.

Here, we show that hormetic HS induced rapid HLH-30 nuclear localization, dephosphorylation of HLH-30 S201, and inhibition of mTORC1 kinase activity. However, neither altering HLH-30 S201 phosphorylation status nor genetic manipulation of mTORC1 signaling prevented HS-induced HLH-30 nuclear localization or long-term hormetic protection. HS also activated AAK-2 kinase activity, and *aak-2* was required for hormetic protection. However, neither *aak-2* deletion nor deletion of the HLH-30 C-terminal region containing residues corresponding to the AMPK-regulated serine cluster in mammalian TFEB prevented HS-induced HLH-30 nuclear localization. *aak-2* was also dispensable for HS-induced autophagosome formation. Together, these findings indicate that although hormetic HS engages canonical nutrient- and energy-sensing pathways, mTORC1- and AMPK-dependent mechanisms do not account for HS-induced HLH-30 nuclear localization, suggesting that additional mechanisms regulate HLH-30 during heat stress in *C. elegans*.

## RESULTS

### HLH-30 S201 phosphorylation is dispensable for heat shock-induced nuclear localization and hormetic benefits

A brief, sub-lethal heat shock (HS) is a potent inducer of HLH-30/TFEB nuclear translocation in young adult *C. elegans* [6]. To better define the dynamics of this response, we first characterized the kinetics of HLH-30 nuclear localization during HS. Under basal conditions, HLH-30::GFP was largely diffuse with no nuclear enrichment (**Fig. 1A**, **B**). HS at 36°C induced HLH-30::GFP nuclear translocation within 15-30 min, with progressively stronger nuclear enrichment over the course of the one-hour HS treatment (**Fig. 1A**, **B**). Maximal nuclear localization was reached within 45 minutes of exposure to 36°C. HLH-30 nuclear localization was quantified both by counting HLH-30::GFP-enriched nuclei per animal and using a semi-quantitative scoring system ranging from 0 (diffuse localization) to 5 (complete nuclear localization) (**Fig. 1B**; **Fig. S1**). The two approaches showed similar kinetics of HS-induced HLH-30 nuclear localization (**Fig. 1A**), supporting use of the 0-5 scoring system for subsequent experiments.

**Figure 1.**
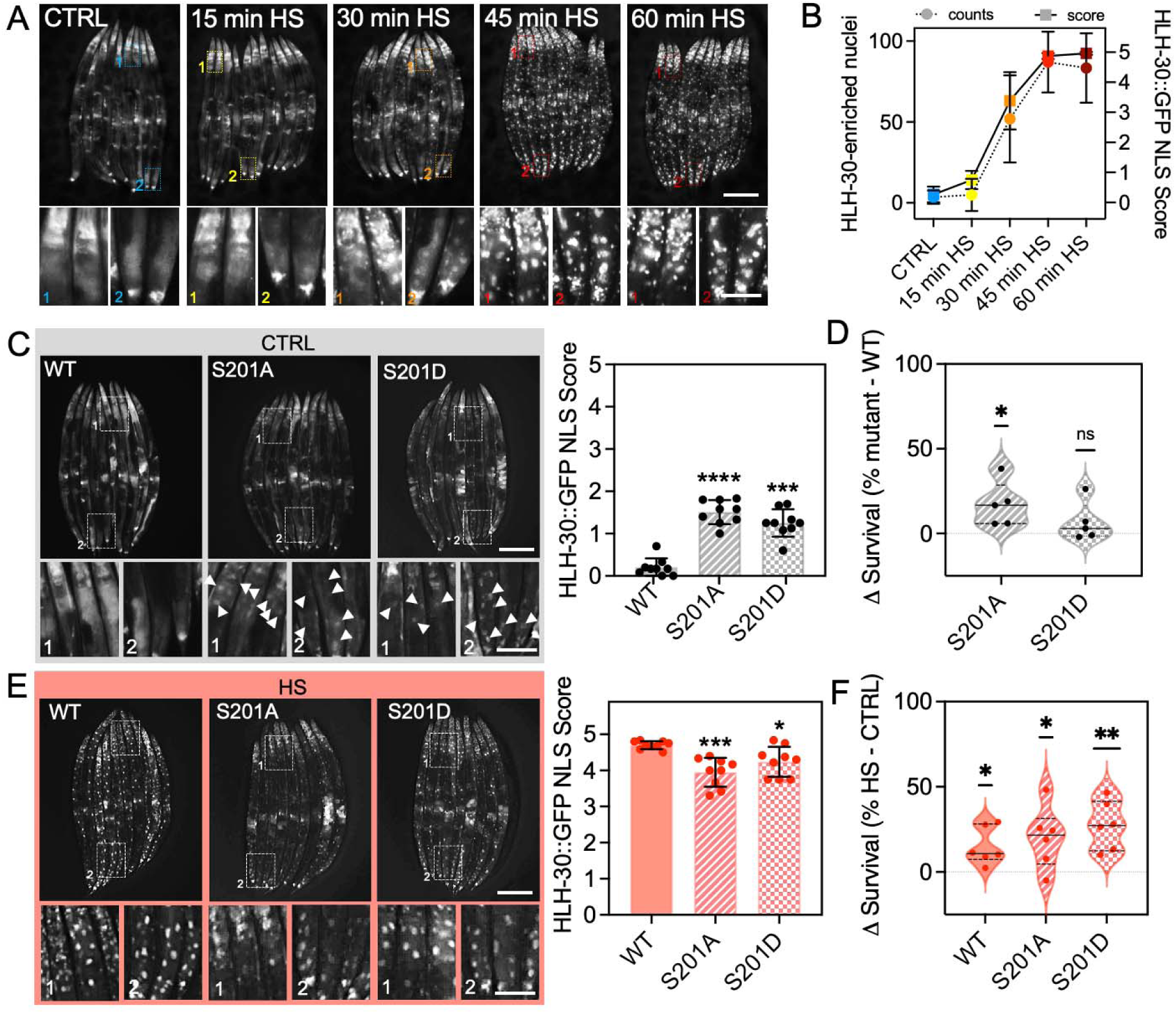
Heat shock induces HLH-30 nuclear localization and hormetic stress resistance independently of S201 phosphorylation. **(A)** Representative fluorescence images of HLH-30::GFP in day 1 adult animals under unstressed conditions (CTRL; 20°C) or following heat shock (HS; 36°C) for the indicated durations. Enlarged regions show representative HLH-30::GFP localization patterns in the head (1) and intestinal (2) regions. **(B)** Quantification of HLH-30::GFP nuclear localization in animals shown in (A) using two approaches: direct counting of HLH-30::GFP-enriched nuclei per animal (left y-axis, circles, dashed line) and a semi-quantitative nuclear localization score ranging from 0 (diffuse localization) to 5 (complete nuclear localization) (right y-axis, squares, solid line; scoring criteria shown in **Fig. S1**). For direct nuclear counts, all individual animals were pooled across experiments and plotted as mean ± SD (N = 54-57 animals/time point). For nuclear localization scoring, a mean score was calculated for each independent experiment from 9-12 animals/time point, and data are shown as the mean ± SD of five independent experiments (n = 5). **(C)** Representative fluorescence images and quantification of HLH-30::GFP nuclear localization in wild-type (WT), phospho-null HLH-30 S201A, and phospho-mimetic HLH-30 S201D animals under unstressed conditions (CTRL; 20°C). Enlarged regions correspond to the boxed head (1) and intestinal (2) regions; arrowheads indicate HLH-30::GFP-enriched nuclei. Nuclear localization was scored from 0 to 5 as described in (B). Data are mean ± SD of n = 9 independent experiments (5-12 animals/condition/experiment). ****P < 0.0001, ***P < 0.001 versus WT by one-way ANOVA with Dunnett’s multiple-comparisons test. **(D)** Difference in survival following prolonged heat stress on day 3 of adulthood between each S201 mutant and WT (Δ survival = % survival mutant − % survival WT). Each point represents one independent experiment (n = 5); solid and dashed lines indicate the median and quartiles, respectively. *P < 0.05; ns, not significant, by two-tailed one-sample t-test against zero. **(E)** Representative fluorescence images and quantification of HLH-30::GFP nuclear localization in wild-type (WT), phospho-null HLH-30 S201A, and phospho-mimetic HLH-30 S201D animals after 1 h heat shock (HS; 36°C). Enlarged regions correspond to the boxed head (1) and intestinal (2) regions; arrowheads indicate HLH-30::GFP-enriched nuclei. Nuclear localization was scored from 0 to 5 as described in (B). Data are mean ± SD of n = 9 independent experiments (5–12 animals/condition/experiment). ***P < 0.001, *P < 0.05 versus WT by one- way ANOVA with Dunnett’s multiple-comparisons test. **(F)** Hormetic benefit in WT and S201 mutant animals, measured as the difference in survival following prolonged heat stress on day 3 of adulthood between animals exposed to hormetic HS on day 1 and their corresponding CTRL animals (Δ survival = % survival hormetic HS − % survival CTRL). Each point represents one independent experiment (n = 5); solid and dashed lines indicate the median and quartiles, respectively. **P < 0.01, *P < 0.05 by two-tailed one- sample t-test against zero. Scale bar: 200 µm; scale bar in magnified ROIs: 50 µm.

To investigate how HS activates HLH-30 nuclear translocation, we asked whether hormetic HS alters post-translational modifications (PTMs) of HLH-30. Although sequence coverage of HLH-30 was limited, mass spectrometry identified phosphorylation of HLH-30 S201 under control conditions that was no longer detected following a 1h hormetic HS on day 1 of adulthood (**Fig. S2**). HLH-30 S201 corresponds to S211 in human TFEB, a conserved mTORC1-regulated site that plays a major role in TFEB cytoplasmic retention [12, 13] (**Fig. S2**). The loss of detectable HLH-30 S201 phosphorylation following HS therefore suggests that HS-induced HLH-30 nuclear translocation could involve regulation via this conserved phosphosite.

To determine whether HLH-30 S201 regulates nuclear translocation in response to HS, we generated transgenic strains expressing phospho-null (S201A) and phospho-mimetic (S201D) HLH-30. We first examined HLH-30 localization under basal conditions and found that strains expressing either HLH-30 S201A or S201D exhibited modestly increased nuclear localization relative to wild type (**Fig. 1C**), consistent with previous findings using endogenous S201 substitutions [15]. We next asked whether these modest increases in basal HLH-30 nuclear localization translated into improved stress resistance. Whereas the phospho-null HLH-30 S201A mutation increased basal thermotolerance, the HLH-30 S201D mutation did not (**Fig. 1D**), indicating that modest increases in basal HLH-30 nuclear localization do not consistently predict stress resistance.

To test whether HLH-30 S201 phosphorylation is required during hormetic HS, we examined HLH-30 nuclear localization and thermotolerance in animals expressing HLH-30 S201A or S201D. HS induced robust HLH-30 nuclear localization in both HLH-30 S201A- and S201D- expressing animals, although nuclear localization was modestly reduced relative to wild type (**Fig. 1E**). Hormetic HS increased thermotolerance in both HLH-30 S201A-, and S201D- expressing animals, comparable to wild-type animals (**Fig. 1F**). Together, these findings demonstrate that although substitutions at HLH-30 S201 altered basal HLH-30 localization, neither HLH-30 S201A nor S201D prevented HS-induced nuclear translocation or hormetic stress resistance.

### Hormetic heat shock and mTORC1 inhibition promote stress resistance through distinct mechanisms

In parallel with testing the requirement for HLH-30 S201 phosphorylation, we asked whether hormetic HS alters mTORC1 activity and whether manipulating mTORC1 signaling modulates the hormetic benefits of HS via HLH-30. We first examined whether hormetic HS inhibits mTORC1 by measuring phosphorylation of the mTORC1 substrate RSKS-1/S6K [27]. HS caused a reduction in pRSKS-1, consistent with inhibition of mTORC1 kinase activity (**Fig. 2A**). To determine whether mTORC1 inhibition is sufficient to promote HLH-30 nuclear localization, we depleted *let-363* by RNAi. *let-363* RNAi induced modest HLH-30 nuclear localization (**Fig. 2B**), whereas HS induced robust nuclear accumulation (**Fig. 2B**), and both *let-363* RNAi and hormetic HS increased thermotolerance (**Fig. 2C**). Together, these findings suggest that HLH-30 nuclear localization and stress resistance can be uncoupled.

**Figure 2.**
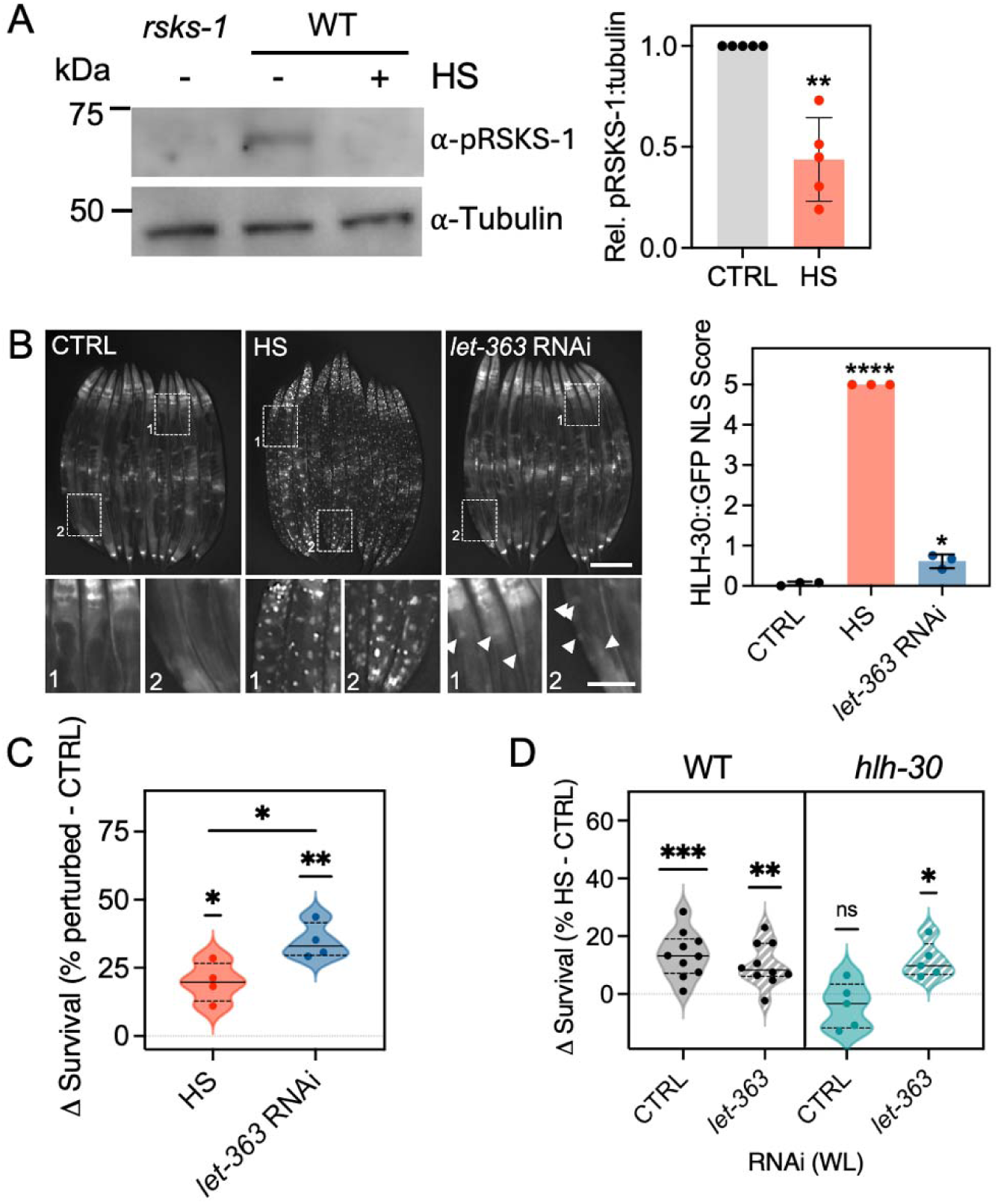
Heat shock-induced HLH-30 nuclear localization and hormetic stress resistance are independent of mTORC1 inhibition. **(A)** Representative Western blot and quantification of phospho (p) RSKS-1(Thr404) in day 1 adult wild-type (WT) animals under unstressed conditions (CTRL; 20°C) or following heat shock (HS; 1 h, 36°C). *rsks-1* null mutants were used as a negative control for antibody specificity. pRSKS-1 levels were normalized to α-tubulin and expressed relative to CTRL. Data are mean ± SD of n = 5 independent experiments. **P < 0.01 by two-tailed t-test. **(B)** Representative fluorescence images and quantification of HLH-30::GFP nuclear localization in WT animals under unstressed conditions (CTRL), following HS (1 h, 36°C), or following *let-363* RNAi from hatching under unstressed conditions. Enlarged regions correspond to the boxed head (1) and intestinal (2) regions; arrowheads indicate HLH-30::GFP-enriched nuclei. Nuclear localization was scored from 0 to 5. Data are mean ± SD of n = 3 independent experiments (12 animals/condition/experiment). ****P < 0.0001, *P < 0.05 versus CTRL by one- way ANOVA with Dunnett’s multiple-comparisons test. Scale bar: 200 µm; scale bar in magnified ROIs: 50 µm. **(C)** Difference in survival following prolonged heat stress on day 3 of adulthood in WT animals exposed to hormetic HS (1 h, 36°C) on day 1 or treated with *let-363* RNAi from hatching, relative to their corresponding controls (Δ survival = % survival perturbed − % survival CTRL). Each point represents one independent experiment (n = 4); solid and dashed lines indicate the median and quartiles, respectively. *P < 0.05, **P < 0.01 by two-tailed one-sample t-test against zero. The comparison *P < 0.05 between HS and *let-363* RNAi was performed by paired, two- tailed t-test. **(D)** Difference in survival following prolonged heat stress on day 3 of adulthood in WT and *hlh- 30* mutant animals treated with control or *let-363* RNAi from hatching. Within each genotype and RNAi condition, animals were either maintained at 20°C or exposed to hormetic HS (1 h, 36°C) on day 1 of adulthood. Δ survival was calculated as the difference between animals receiving hormetic HS and their corresponding unstressed controls (% survival HS − % survival CTRL). Each point represents one independent experiment (WT, n = 10; *hlh-30*, n = 5); solid and dashed lines indicate the median and quartiles, respectively. ***P < 0.001, **P < 0.01, *P < 0.05; ns, not significant, by two-tailed one-sample t-test against zero.

To further distinguish the effects of mTORC1 signaling from those of hormetic HS, we tested whether constitutive mTORC1 activation would suppress HS-induced HLH-30 nuclear localization and stress resistance. To test this, we used a gain-of-function mutation of RAGA-1, an upstream activator of mTORC1, *raga-1(viz128[Q63L])* (*raga-1(gf)*) [28], which has previously been shown to increase mTORC1 activity, as evidenced by increased RSKS-1 phosphorylation [29]. However, HS robustly reduced RSKS-1 phosphorylation in *raga-1(gf)* animals (**Fig. S3A**), indicating that HS can inhibit mTORC1 despite constitutive RAGA-1 activation. Consistent with this, HS-induced HLH-30 nuclear localization remained robust in *raga-1(gf)* mutants (**Fig. S3B**). Thus, RAGA-1-mediated activation was unable to maintain mTORC1 activity during HS, precluding a direct test of whether sustained mTORC1 activity is sufficient to block HS-induced HLH-30 nuclear translocation.

To determine whether HS and mTORC1 inhibition provide protection through the same mechanism, we combined hormetic HS with *let-363* RNAi. In wild-type animals, hormetic HS increased thermotolerance under both control and *let-363* RNAi conditions (**Fig. 2D**), indicating that hormetic HS provides additional protection when mTORC1 is already reduced. Thus, mTORC1 inhibition alone does not account for HS-induced stress resistance.

Since *hlh-30* is required for lifespan extension upon mTORC1 inhibition [8], we asked whether the protective effect of mTORC1 inhibition in our thermotolerance paradigm similarly depended on *hlh-30*. As expected, *hlh-30* mutants failed to benefit from hormetic HS under control RNAi conditions (**Fig. 2D**). Strikingly, however, HS increased thermotolerance in *hlh-30* mutants when mTORC1 was inhibited by *let-363* RNAi (**Fig. 2D**). Thus, while HLH-30 is required for hormetic protection under normal mTORC1 conditions, reducing mTORC1 activity reveals an HLH-30-independent component of HS-induced protection.

Together, these findings demonstrate that although hormetic HS inhibits mTORC1, mTORC1 inhibition alone does not recapitulate the HS response. Instead, HS and mTORC1 inhibition engage distinct but interacting protective mechanisms.

### AMPK is required for hormetic protection but dispensable for HLH-30 nuclear localization and autophagosome formation

AMPK is an energy-sensing kinase implicated in TFEB regulation and autophagy induction [19–21, 23, 30]. We therefore examined whether AMPK contributes to HS-induced HLH-30 regulation and hormetic benefits. Hormetic HS has been shown to reduce cellular energy levels and require *aak-2*/AMPK for lifespan extension in *C. elegans* [25]. Consistent with AAK-2 activation under these conditions, hormetic HS increased phosphorylation of AAK-2 at Thr243, corresponding to the conserved activating Thr172 phosphorylation site in mammalian AMPK [31] (**Fig. 3A**). Moreover, loss of *aak-2* abolished the increase in thermotolerance conferred by hormetic HS (**Fig. 3B**), confirming a requirement for *aak-2* in hormetic stress resistance.

**Figure 3.**
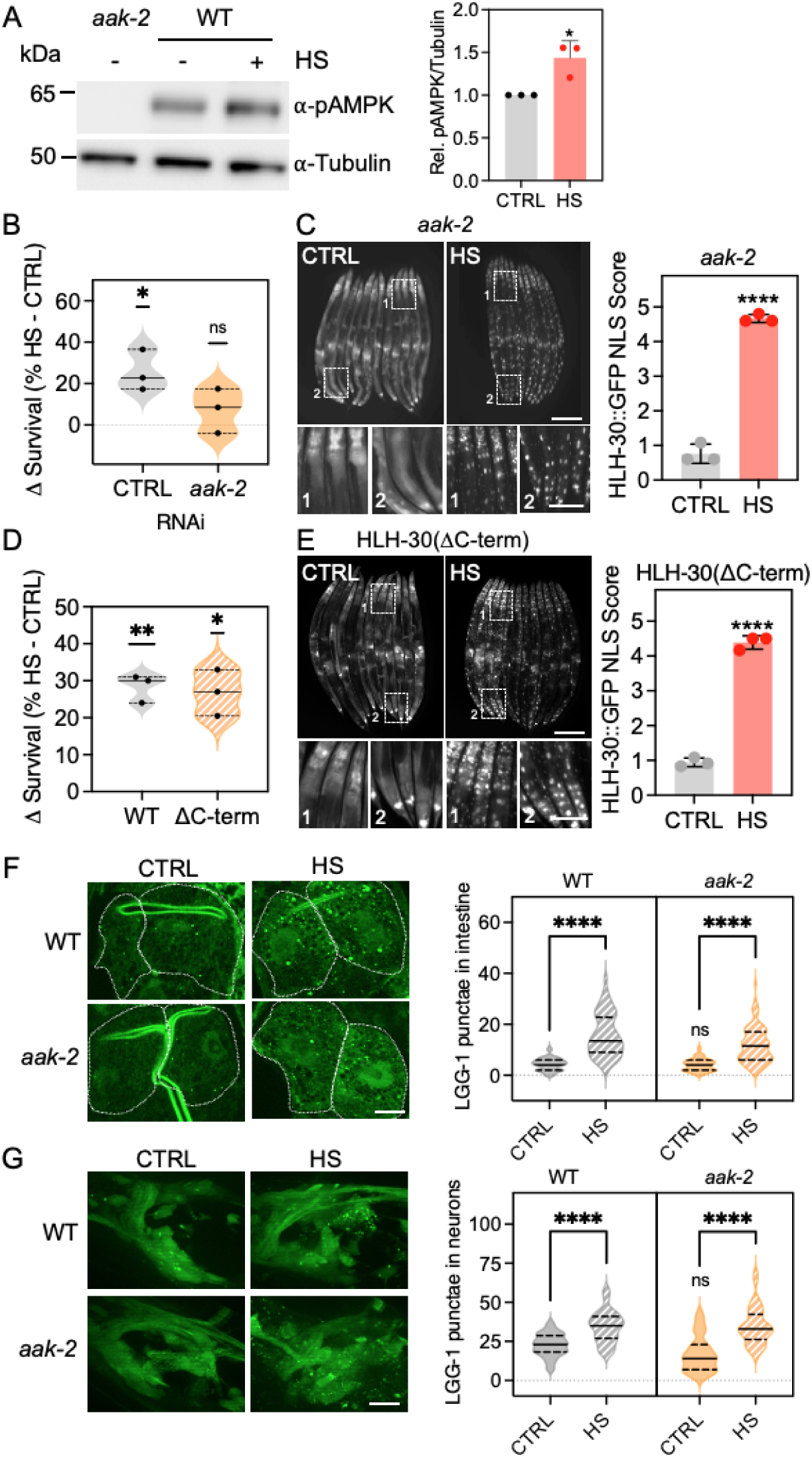
AMPK is activated by heat shock, required for hormetic stress resistance but dispensable for heat shock-induced HLH-30 nuclear localization and autophagosome formation. (A) Representative Western blot and quantification of phospho (p) AAK-2(Thr243) in day 1 adult *aak-2(ok524)* and wild-type (WT) animals under unstressed conditions (CTRL; 20°C) or following heat shock (HS; 1 h, 36°C). *aak-2(ok524)* mutants were used as a negative control for antibody specificity. pAAK-2 levels were normalized to α-tubulin and expressed relative to CTRL. Data are mean ± SD of n = 3 independent experiments. *P < 0.05 by unpaired, two-tailed t-test. (B) Difference in survival following prolonged heat stress on day 3 of adulthood in animals treated with control or *aak-2* RNAi from adulthood on. Within each RNAi condition, animals were either maintained at 20°C or exposed to hormetic HS (1 h, 36°C) on day 1 of adulthood. Δ survival was calculated as the difference between animals receiving hormetic HS and their corresponding unstressed controls (% survival HS − % survival CTRL). Each point represents one independent experiment (n = 3); solid and dashed lines indicate the median and quartiles, respectively. *P < 0.05; ns, not significant, by two-tailed one-sample t-test against zero. (C) Representative fluorescence images and quantification of HLH-30::GFP nuclear localization in *aak-2* mutant animals under unstressed conditions (CTRL) or following HS (1 h, 36°C). Enlarged regions correspond to the boxed head (1) and intestinal (2) regions. Nuclear localization was scored from 0 to 5. Data are mean ± SD of n = 3 independent experiments (N = 12 animals/experiment). ****P < 0.0001 by unpaired, two-tailed t-test. Scale bar: 200 μm; scale bar in magnified ROIs: 50 µm. (D) Difference in survival following prolonged heat stress on day 3 of adulthood in WT and HLH- 30(ΔC-term) animals. Within each genotype, animals were either maintained at 20°C or exposed to hormetic HS (1 h, 36°C) on day 1 of adulthood. Δ survival was calculated as the difference between animals receiving hormetic HS and their corresponding unstressed controls (% survival HS − % survival CTRL). Each point represents one independent experiment (n = 3); solid and dashed lines indicate the median and quartiles, respectively. **P < 0.01, *P < 0.05 by two-tailed one-sample t-test against zero. (E) Representative fluorescence images and quantification of HLH-30::GFP nuclear localization in HLH-30(ΔC-term) animals under unstressed conditions (CTRL) or following HS (1 h, 36°C). Enlarged regions correspond to the boxed head (1) and intestinal (2) regions. Nuclear localization was scored from 0 to 5. Data are mean ± SD of n = 3 independent experiments (N = 12 animals/experiment). ****P < 0.0001 by unpaired, two-tailed t-test. Scale bar: 200 μm; scale bar in magnified ROIs: 50 µm. (F) Representative fluorescence images and quantification of GFP::LGG-1-positive puncta in the intestine of WT and *aak-2* mutant animals expressing *lgg-1p::gfp::lgg-1* under unstressed conditions (CTRL) or following HS (1 h, 36°C). Dashed outlines indicate the intestinal region used for quantification. n = 3 independent experiments with a total of N = 56-76 cells. Solid and dashed lines indicate the median and quartiles, respectively. ****P < 0.0001; ns, not significant, by two-way ANOVA with Sidaks multiple-comparison tests. (G) Representative fluorescence images and quantification of GFP::LGG-1-positive puncta in neurons of WT and *aak-2* mutant animals expressing *rgef-1p::gfp::lgg-1* under unstressed conditions (CTRL) or following HS (1 h, 36°C). n = 3 independent experiments with a total of N = 31-36 animals. Solid and dashed lines indicate the median and quartiles, respectively. ****P < 0.0001; ns, not significant, by Two-way ANOVA with Sidaks multiple comparison tests.

AMPK-dependent regulation of mammalian TFEB has been linked to a conserved C-terminal serine cluster comprising TFEB S466/S467/S469 [20]. The corresponding residues are conserved in HLH-30 as Ser465/476/479 (**Fig. S2A**). Despite the requirement for *aak-2* in hormesis, HS induced robust HLH-30 nuclear localization in *aak-2* mutants (**Fig. 3C**). Similarly, deletion of the HLH-30 C-terminal region containing the conserved serine cluster did not impair thermotolerance (**Fig. 3D**) or HS-induced nuclear localization (**Fig. 3E**). Thus, neither *aak-2* nor the HLH-30 C-terminal region are required for HLH-30 nuclear localization during hormetic HS.

Since *aak-2* was required for the benefits of a hormetic HS (**Fig. 3B)** [25], but not HLH-30 nuclear localization (**Fig. 3C**), we examined whether *aak-2* was required for HS-induced autophagy, another essential component of the hormetic response [5, 6]. Hormetic HS induces autophagy, as previously established using autophagic flux assays [5, 6]. To assess the autophagy response in *aak-2* mutants after hormetic HS, we quantified LGG-1/Atg8 puncta, which mark autophagosomes. HS-induced accumulation of LGG-1/Atg8 puncta was preserved in *aak-2* mutants in both the intestine and neurons (**Fig. 3F-G**). Together, these findings demonstrate that although AAK-2 is activated by hormetic HS and required for the HS-induced hormetic benefits, it is dispensable for HS-induced HLH-30 nuclear localization and autophagosome formation.

## DISCUSSION

Hormetic HS induced rapid HLH-30/TFEB nuclear localization together with several signaling events associated with canonical TFEB regulation, including mTORC1 inhibition, loss of detectable phosphorylation at the conserved HLH-30 S201 residue, and activation of AAK- 2/AMPK. Surprisingly, however, these events did not explain HLH-30 nuclear translocation during HS. Substitution of HLH-30 S201 did not prevent HS-induced HLH-30 nuclear localization or hormetic protection, while *aak-2* was required for hormetic protection but dispensable for HLH-30 nuclear localization and autophagosome formation (**Fig. 4**). Together, these findings suggest that the canonical relationships between nutrient-sensing pathways and HLH-30 established under basal conditions are altered during heat stress, with mTORC1, AMPK, and HLH-30 making distinct and context-dependent contributions to hormetic protection (**Fig. 4**).

**Figure 4.**
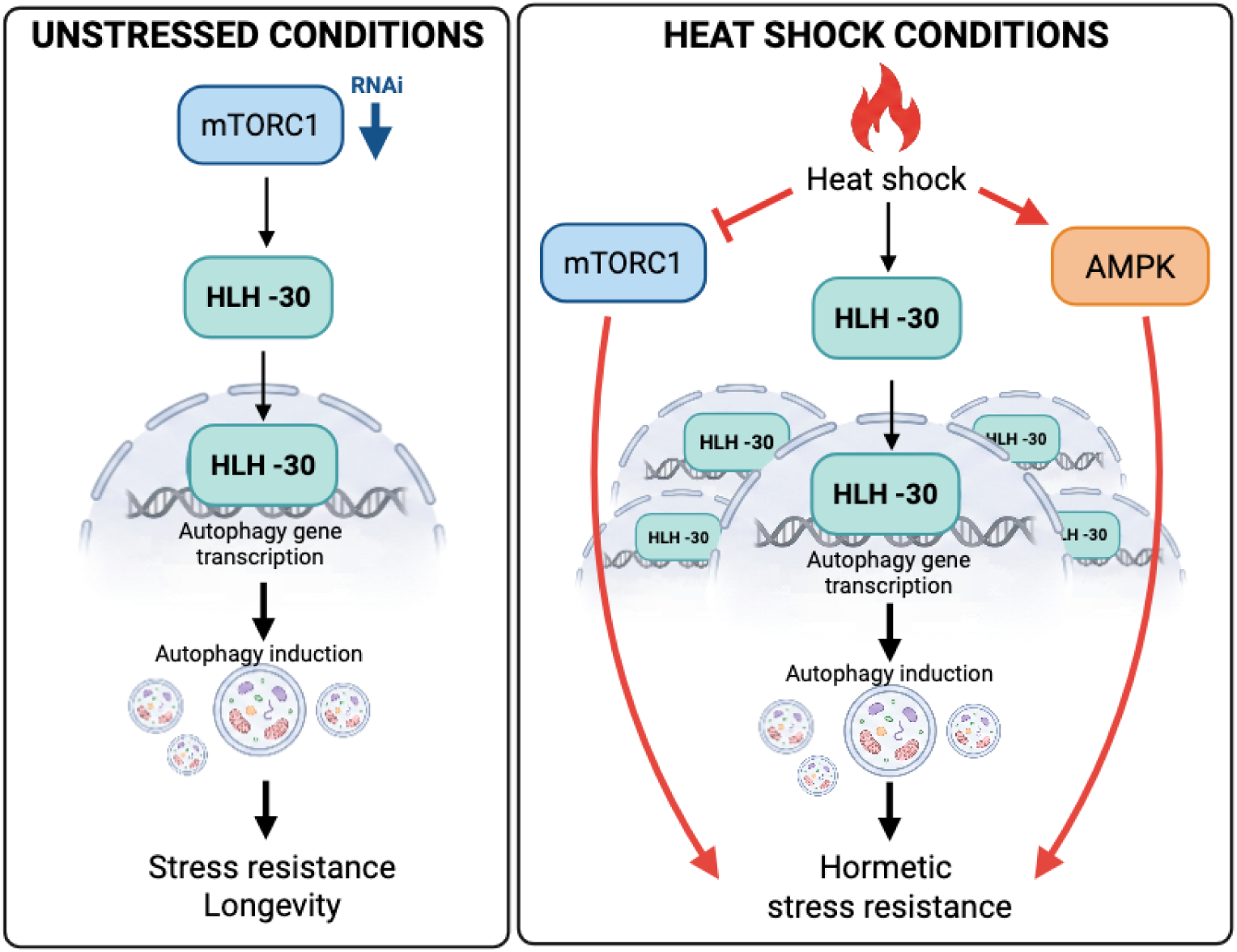
Model of HLH-30 regulation under unstressed and hormetic heat shock conditions. Under unstressed conditions, inhibition of mTORC1 by *let-363* RNAi promotes mild HLH-30 nuclear localization and HLH-30-dependent transcription of autophagy genes, autophagy induction, stress resistance, and longevity. During hormetic heat shock (HS), HLH-30 robustly translocates to the nucleus and promotes autophagy gene transcription and autophagy, contributing to hormetic stress resistance. At the same time, HS inhibits mTORC1 and activates AMPK; however, neither canonical mTORC1-dependent regulation nor AMPK is required for HS-induced HLH-30 nuclear localization. Instead, reduced mTORC1 activity and AMPK make additional contributions to hormetic stress resistance. Specifically, inhibition of mTORC1 reveals an HLH-30-independent component of HS-induced stress resistance, whereas AMPK is required for hormetic protection despite being dispensable for HS-induced HLH-30 nuclear localization and autophagosome formation. Red arrows and inhibitory connections indicate relationships identified in this study; black arrows indicate previously established relationships.

Previous work showed that endogenous substitution of HLH-30 S201 with either alanine or aspartate increases basal nuclear localization without inducing a robust transcriptional response [15]. Consistent with these findings, we observed similarly increased basal nuclear localization in our transgenic strains expressing HLH-30 S201A and S201D. However, despite their similar effects on basal nuclear localization, only HLH-30 S201A increased basal thermotolerance. Moreover, neither substitution prevented the substantially stronger HLH-30 nuclear localization induced by HS, and both mutants retained hormetic protection. Thus, S201-dependent regulation represents one layer of HLH-30 control, while additional mechanisms engaged during stress appear to determine its transcriptional and physiological outputs.

Despite being insufficient to explain HLH-30 nuclear localization, mTORC1 signaling was affected by HS. HS reduced RSKS-1 phosphorylation even in *raga-1(gf)* animals, indicating that heat stress can inhibit mTORC1 despite RAGA-1-dependent activation. HS may therefore inhibit mTORC1 downstream of, or in parallel to, Rag-dependent nutrient signaling. In mammalian cells, heat stress has been reported to inhibit mTORC1 through c-Jun N-terminal kinase (JNK)- dependent phosphorylation of mTOR and Raptor, while heat shock factor 1 (HSF1) can modulate this response through sequestration of JNK [32]. A related stress-responsive mechanism could explain why constitutive RAGA-1 activation was unable to maintain mTORC1 activity during HS. At the organismal level, however, HS and mTORC1 inhibition were not equivalent. HS provided additional protection when mTORC1 was reduced, and, strikingly, HS increased stress resistance in *hlh-30* mutants under *let-363* RNAi conditions despite the requirement for HLH-30 under control conditions. These findings suggest that HS engages both HLH-30-dependent and HLH-30-independent protective mechanisms (**Fig. 4**). mTORC1 inhibition may shift animals into a physiological state in which other HS-induced protective mechanisms become sufficient to compensate for the absence of HLH-30. In either case, the requirement for HLH-30 is conditional on mTORC1 signaling state rather than fixed, highlighting a fundamental difference between mTORC1-HLH-30 interactions under basal and heat-stress conditions.

AAK-2/AMPK revealed a related separation between pathway activation and functional output. HS activated AAK-2, and loss of *aak-2* abolished hormetic protection, consistent with an essential role for AMPK in the response. However, neither loss of *aak-2* nor deletion of the HLH- 30 C-terminal region containing the conserved TFEB serine cluster impaired HS-induced HLH-30 nuclear localization, and the C-terminal deletion did not impair hormetic protection. The relationship between AMPK and the conserved TFEB C-terminal serine cluster remains unresolved. AMPK has been reported to promote TFEB transcriptional activity through phosphorylation of S466/S467/S469 without affecting nuclear localization [20], whereas more recent work found that AMPK activation instead promotes dephosphorylation of these residues and that the serine cluster is a poor direct AMPK substrate [21]. Despite these mechanistic differences, both studies support a role for AMPK in regulating TFEB transcriptional activity. Our findings demonstrate that neither *aak-2* nor the corresponding HLH-30 C-terminal region is required for HS-induced HLH-30 nuclear localization, but do not address whether AAK-2 regulates HLH-30 transcriptional activity after nuclear translocation. Moreover, HS-induced LGG-1/Atg8 puncta formation was preserved in *aak-2* mutants in both the intestine and neurons. This contrasts with the ability of constitutively active AAK-2 to increase LGG-1-positive puncta through an UNC-51-dependent mechanism in *C. elegans* hypodermal seam cells [22], suggesting that the contribution of AAK-2 to autophagosome formation depends on tissue and physiological context. AAK-2 may instead promote hormetic protection through downstream mechanisms distinct from those regulating HLH-30 localization or autophagosome formation. Determining which AAK-2-dependent outputs mediate hormetic thermotolerance will therefore be important for understanding how AMPK contributes to the HS response despite being dispensable for HLH-30 nuclear localization and autophagosome formation.

More broadly, our findings demonstrate that HLH-30 nuclear localization does not necessarily predict its physiological output. Across HLH-30 S201 substitution, mTORC1 inhibition, and loss of *aak-2*, HLH-30 nuclear localization and stress resistance could be uncoupled, indicating that nuclear localization represents only one level of HLH-30 regulation. Together, our findings show that hormetic HS does not simply engage the canonical nutrient-sensing mechanisms that regulate HLH-30 under basal conditions. Instead, additional stress- responsive mechanisms appear to drive HLH-30 nuclear localization, while mTORC1, AAK-2, and HLH-30 make distinct and context-dependent contributions to hormetic protection. Defining these stress-specific mechanisms will be important for understanding how HLH-30/TFEB coordinates adaptive responses to different physiological challenges.

## METHODS

### *C. elegans* maintenance and RNA interference

*C. elegans* were maintained at 20°C on nematode growth medium (NGM) plates seeded with *Escherichia coli* OP50 and cultured according to standard protocols [33], except where otherwise stated. All *C. elegans* strains used in this study, including strain names, genotypes, sources, and generation details, are listed in **Supplemental Table 1**. Plasmids used to generate transgenic strains, including their construction and primer information, are described in **Supplemental Table 2**.

For RNAi experiments, animals were fed *E. coli* HT115 expressing double-stranded RNA (dsRNA) targeting the gene of interest. HT115 bacteria carrying RNAi constructs were grown in Luria-Bertani (LB) medium containing 0.1 mg/ml carbenicillin (Carb; BioPioneer). Bacterial cultures (80 µl) were spotted onto 6-cm NGM plates containing 0.1 mg/ml Carb and allowed to grow for 1-2 days. To induce dsRNA expression, 80 µl of a solution containing 0.1 M isopropyl-β-D-thiogalactoside (IPTG; Promega/Gold Biotechnology) and 200 mg/ml Carb was added directly to the bacterial lawn. The *let-363* RNAi clone was provided by Dr. J. Avruch. For *let-363* RNAi experiments, animals were exposed to RNAi from hatching and maintained on *let-363* RNAi throughout life. To confirm RNAi efficacy, eggs from RNAi-treated animals were transferred to *let-363* RNAi plates, where the expected larval-arrest phenotype was verified. The *aak-2* RNAi clone was obtained from the Ahringer library [34]. For *aak-2* RNAi experiments, animals were transferred to *aak-2* RNAi on day 1 of adulthood and maintained on RNAi until experimental endpoint. As control, L4440 empty vector was used in all RNAi experiments, and all RNAi clones were verified by sequencing.

### Fluorescence imaging and quantification

For fluorescence imaging, worms were immobilized in M9 containing 150 µM sodium azide and imaged using a Leica DFC310 FX camera with fluorescence and brightfield illumination. Images were acquired at 16x magnification. Within each experiment, exposure settings were kept constant across all conditions used for quantitative comparisons, and quantification was performed on raw images. HLH-30::GFP nuclear localization was quantified using either direct counting of HLH-30::GFP-enriched nuclei (**Fig. 1B**) and reported as the number of enriched nuclei per animal, or a semi-quantitative nuclear localization score on a scale from 0 to 5 (**Fig. 1B**) for all subsequent experiments. Representative localization patterns are shown in **Fig. S1**. A score of 0 indicated no detectable nuclear enrichment; 1, enrichment in a small number of nuclei at either the anterior or posterior end of the animal; 2, enrichment in nuclei at both ends of the animal; 3, mild nuclear enrichment throughout the body; 4, increased number and intensity of enriched nuclei throughout the body; and 5, strong nuclear enrichment throughout the animal, including large intestinal nuclei and smaller nuclei in the head region. For each experiment, 6-12 animals per condition were scored and the mean nuclear localization score was calculated for each condition. Experiments were performed independently at least three times, with exact sample sizes and numbers of independent experiments indicated in the corresponding figure legends.

### Thermorecovery assays in *C. elegans*

Age-synchronized animals were grown under the indicated experimental conditions and distributed onto four assay plates per condition (25 animals/plate). On day 3 of adulthood, animals were exposed to prolonged heat stress at 36°C for 5-7 h and subsequently returned to 20°C for recovery. Survival was scored the next day. Animals displaying spontaneous movement were scored as alive. Non-moving animals were gently prodded with a platinum wire and scored as dead if they failed to respond to touch. Survival was calculated separately for each plate. The four plate-level values were averaged to obtain one survival value per condition for each independent experiment, and independent experiments were used as the biological replicates for statistical analysis. Thermotolerance benefit was expressed as the change in survival relative to the corresponding control in paired experiments (Δ Survival = % survival experimental − % survival control), including comparisons of hormetic HS, genetic mutations, and RNAi treatments. Independent experiments and sample sizes are indicated in the corresponding figure legends.

### Mass spectrometry analysis of HLH-30 phosphorylation

Mass spectrometry analysis was performed using VZ892/*hlh-30(syb1452[hlh- 30::3xFLAG::eGFP]) IV* animals grown on semi-high growth medium (SHGM) plates. SHGM plates were prepared by combining 100 g agar (BD Difco #DF0145070), 45 g Bacto Peptone (Gibco #211820), 12 g NaCl (Fisher #BP358212) and 4 L dH_2_O and autoclaving. After cooling to approximately 55°C, 4 mL sterile-filtered cholesterol stock (5 mg/mL in ethanol; Specialty Organic Chemicals #103783-604), 4 mL 1 M CaCl_2_ (Acros Organics #AC349615000), 4 mL 1 M MgSO_4_ (Sigma #230391) and 100 mL 1 M potassium phosphate buffer (pH 6.0; 120 g/L KH_₂_PO_₄_ and 21 g/L K_₂_HPO_₄_) were added before plates were poured. Animals were synchronized by bleaching, transferred to fifty 10 cm semi-high-growth medium plates seeded with OP50, and grown at 25°C to day 1 of adulthood. Animals were then divided between control and heat shock conditions and either maintained at 25°C or heat shocked at 36°C for 1 hour. HLH-30::GFP subcellular localization was confirmed by fluorescence microscopy before animals were washed from plates with M9 buffer. Approximately 1.5 mL packed worms per condition was collected and divided among three tubes containing approximately 500 µL worms each. Samples were washed in lysis buffer containing 50 mM HEPES (pH 7.3), 100 mM KCl, 2 mM EDTA, 0.1% NP-40, 10% glycerol, protease inhibitor, and phosphatase inhibitor. Each sample was combined with 500 µL 0.5-mm zirconia beads and disrupted using a FastPrep cell disrupter for 2 min at 50 oscillations/s at 4°C. Lysates were cleared by centrifugation at 500 x *g* for 5 min at 4°C, and protein concentrations were determined using the Pierce 660 nm Protein Assay.

Lysates from the three tubes within each condition were combined, and HLH- 30::3×FLAG::eGFP enrichment was attempted by anti-GFP immunoprecipitation using 1.26 mg and 1.24 mg total protein from control and heat-shocked lysates, respectively. Although enrichment was inefficient, the resulting bead-bound fractions were washed with 50 mM ammonium bicarbonate and submitted to the Sanford Burnham Prebys Proteomics Core for mass spectrometry analysis. HLH-30-derived peptides were detected in both conditions. At the Proteomics Core, proteins were denatured in 8 M urea and 50 mM ammonium bicarbonate, reduced with TCEP, alkylated with iodoacetamide, and diluted to 1 M urea before overnight digestion with mass-spectrometry-grade Trypsin/Lys-C. Digested peptides were acidified and desalted using C18 cartridges on an Agilent AssayMap BRAVO system. Peptides were analyzed by LC-MS/MS using a Proxeon EASY-nanoLC coupled to a Q Exactive Plus mass spectrometer. Spectra were searched using MaxQuant against the *C. elegans* UniProt database, with phosphorylation of serine, threonine, and tyrosine specified as variable modifications and a 1% target-decoy false discovery rate for peptide-spectrum matches and protein identification.

### Western blot analysis

Age-synchronized *C. elegans* were collected on day 1 of adulthood by hand-picking 60-100 animals per sample for phospho-RSKS-1/S6K Western blots or 60 animals per sample for phospho-AAK-2/AMPK Western blots into microcentrifuge tubes containing 30 µL M9 buffer (0.6% sodium phosphate dibasic, 0.3% potassium dihydrogen phosphate, 0.5% sodium chloride, 0.025% magnesium sulfate heptahydrate). Animals were washed 3-5 times with an additional 30 µL M9 buffer per wash to remove residual OP50 bacteria. Following the final wash, the remaining M9 volume was adjusted to approximately 10 µL, and 2 µL 6X Laemmli sample buffer (Thermo Scientific, #J61337.AC) was added. Samples were flash-frozen in liquid nitrogen and lysed by two freeze-thaw cycles, with brief centrifugation following each thaw, and subsequently heated at 95°C for 10 min. Proteins were separated by electrophoresis on 4-12% Bis-Tris gels (NuPAGE) in MOPS running buffer (NuPAGE, #NP0001) at 110 V and transferred to PVDF membranes (Millipore, #IPVH85R) using a Novex Mini Cell system (Invitrogen, #EI0001) at 30 V for 1.5 h on ice. Membranes were blocked in 5% milk in Tris-buffered saline containing 0.05% Tween-20 (TBS-T) for 30 min. RSKS-1 phosphorylation was detected using an antibody recognizing the conserved mTORC1-dependent Thr389 phosphorylation site of mammalian p70 S6 kinase, previously used to detect phosphorylated RSKS-1 in *C. elegans* [27, 29]. AAK-2 phosphorylation at Thr243, corresponding to the conserved activating Thr172 phosphorylation site of mammalian AMPK[31], was detected using anti-phospho-AMPKα (Thr172) antibody (Cell Signaling Technology, #2535), diluted 1:1,000 in 1% milk in TBS-T[35]. Membranes were incubated overnight at 4°C with anti-phospho-p70 S6 kinase (Thr389) antibody (Cell Signaling Technology, #9206) diluted 1:1,000 in 1% milk in TBS-T. Membranes were washed three times for 10 minutes in TBS-T and incubated with HRP-conjugated anti-mouse IgG (Cell Signaling Technology, #7076) diluted 1:10,000 in 1% milk in TBS-T for 2 h at room temperature. Membranes were subsequently washed two times for 10 minutes in TBS-T, developed using SuperSignal chemiluminescent substrate (Thermo Scientific, #34577 or #34095), and imaged using a Bio-Rad ChemiDoc imaging system. For loading normalization, membranes were stripped for 4 min using stripping buffer (Thermo Scientific, #46430), re-blocked in 5% milk in TBS-T, and incubated overnight at 4°C with anti-α-tubulin antibody (Cell Signaling Technology, #2144) diluted 1:1,000 in 1% milk in TBS-T. Following washing, membranes were incubated with HRP-conjugated anti-rabbit IgG (Cell Signaling Technology, #7074) diluted 1:10,000 in 1% milk in TBS-T for 2 h at room temperature, washed, developed, and imaged as described above. Band intensities were quantified using Image Lab v6.1.0 build 7 (Bio-Rad Laboratories, Inc.). For each sample, pRSKS-1 band intensity was normalized to the corresponding α-tubulin band intensity. Normalized pRSKS-1 levels were then expressed relative to the untreated wild-type control within each independent experiment. Biological replicate numbers and statistical analyses are provided in the corresponding figure legends and Statistical Analysis section.

### Autophagy measurements

Autophagy was assessed by quantification of GFP::LGG-1/Atg8-positive puncta [36, 37]. For imaging and puncta quantification, animals were mounted on 2% agarose pads in M9 containing 0.1% NaN_₃_. GFP::LGG-1-positive puncta were imaged at x1,000 magnification using a Zeiss Imager Z1 equipped with an ApoTome.2 and a Hamamatsu ORCA-Flash4LT camera using ZEN 2.3 software. For intestinal measurements, animals expressed *lgg-1p::gfp::lgg-1* [38]. Animals were exposed to hormetic heat shock (HS; 1 h at 36°C) on day 1 of adulthood followed by 4 h of recovery at 20°C, or maintained at the corresponding control temperature. GFP::LGG-1-positive puncta were counted manually in the 2-3 most proximal intestinal cells using a single focal plane containing the intestinal nuclei. For neuronal measurements, animals expressed *rgef- 1p::gfp::lgg-1* [39]. Animals were exposed to hormetic HS (1 h at 36°C) on day 1 of adulthood followed by 2 h of recovery at 20°C, or maintained at the corresponding control temperature. GFP::LGG-1-positive puncta were counted manually in nerve-ring neurons from maximum-intensity projections of Z-stacks comprising 12-15 optical sections acquired at 1.0-µm intervals.

### Quantification and statistical analysis

Statistical analyses and graph generation were performed using GraphPad Prism version 10.6.0. Statistical tests, biological replicate numbers (*n*), sample sizes (*N*), and measures of central tendency and variability are specified in the corresponding figure legends. Unless otherwise indicated, independent experiments were treated as biological replicates. Statistical significance was defined as *P* < 0.05.

## ACKNOWLEDGMENTS

This work was supported by a Conrad Prebys Foundation Predoctoral Fellowship (TMM), NIH-NIA R01-AG083373 (CK), Funding for core facilities: NCI Cancer Center Support Grant P30 CA030199. Some strains were provided by the CGC, which is funded by NIH Office of Research Infrastructure Programs (P40 OD010440). We thank Dr. Antonio Miranda-Vizuete for generously sharing *C. elegans* strains before publication.

## AUTHOR CONTRIBUTIONS

Tatiana M. Moreno: Conceptualization, Investigation, Methodology, Formal Analysis, Visualization, Writing – Review & Editing.

Michelle E. Brown: Conceptualization, Investigation, Methodology, Formal Analysis, Visualization, Writing – Review & Editing.

Caitlin M. Lange: Investigation, Formal Analysis.

Gavin McLaren: Investigation.

Diego A. Hernandez-Urbina: Conceptualization, Writing – Review & Editing. Cheng-Ju Kuo: Conceptualization, Writing – Review & Editing.

Caroline Kumsta: Conceptualization, Resources, Supervision, Project Administration, Funding Acquisition, Writing – Original Draft, Writing – Review & Editing.

## DECLARATION OF GENERATIVE AI AND AI-ASSISTED TECHNOLOGIES

During the preparation of this work, the authors used ChatGPT v5.2 to improve grammar and clarity. After using this tool, the authors reviewed and edited the content as needed and take full responsibility for the content of the publication.

---

**Supplemental Figure 1.**
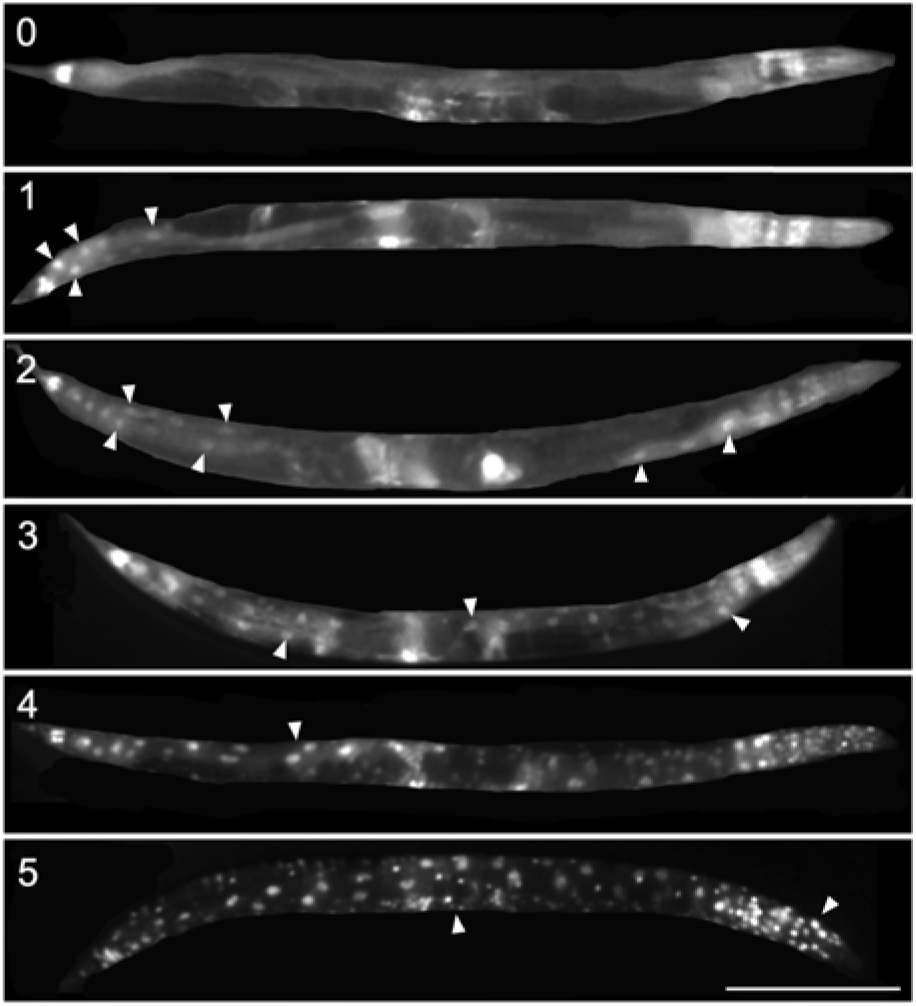
H**L**H**-30::GFP nuclear localization signal scoring scale.** Representative images of HLH-30::GFP nuclear enrichment levels on a scale of 0 to 5. 0, no enriched nuclei; 1, some enriched nuclei in either proximal or distal ends of body; 2, enriched nuclei in both proximal and distal ends of body; 3, mild enrichment throughout the length of the body; 4, increased brightness and number of enriched nuclei throughout the length of the body; 5, complete and bright full body nuclear enrichment in both large intestinal nuclei and small nuclei in the head region. Arrowheads point to progressively greater nuclear enrichment in each worm. Scale bar: 200 µm

**Supplemental Figure 2.**
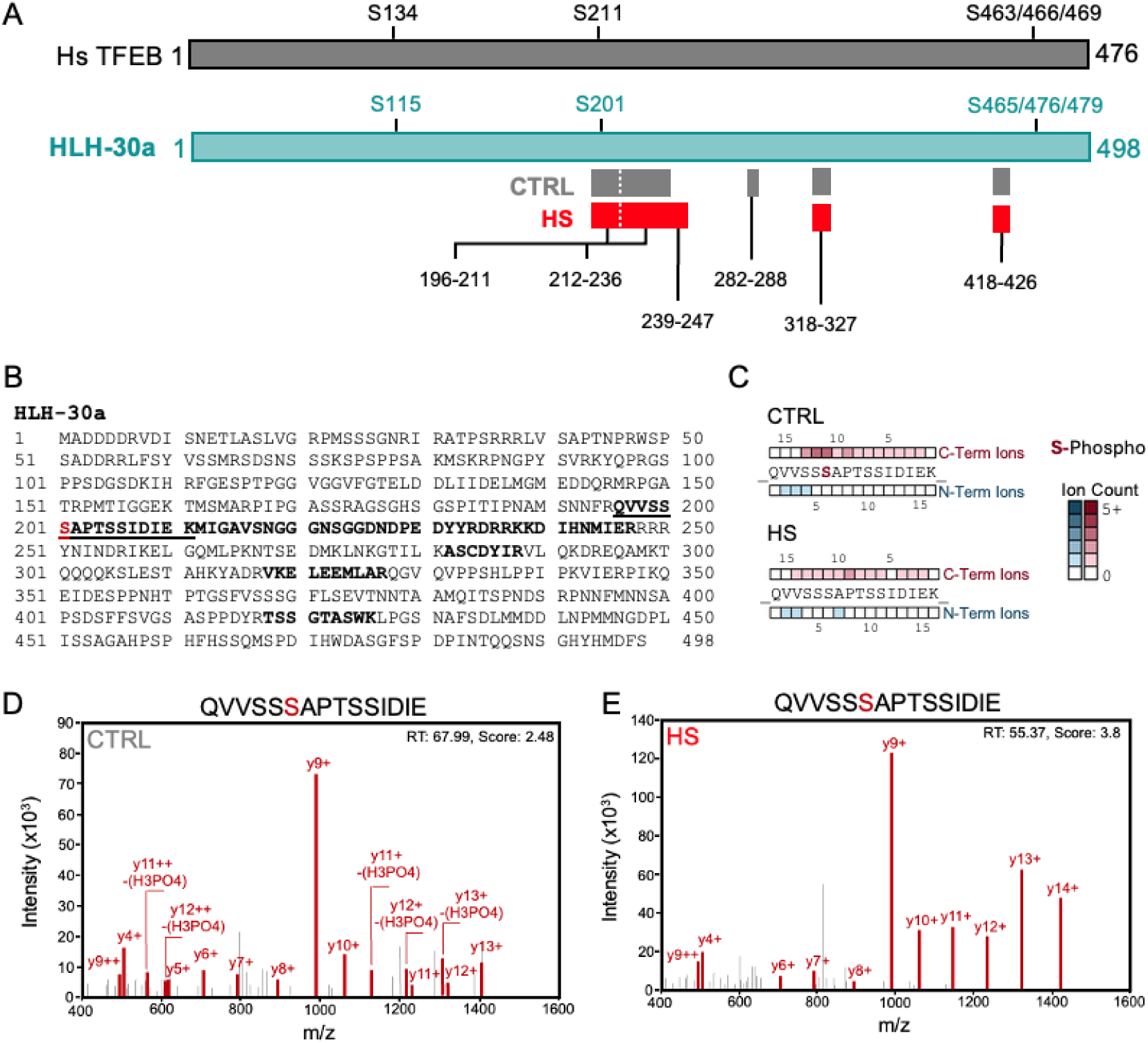
I**d**entification **of heat shock-dependent dephosphorylation of HLH-30 Ser201 by mass spectrometry. (A)** Schematic alignment of human TFEB (Hs TFEB) and *C. elegans* HLH-30 isoform a. Conserved serine residues are indicated, including HLH-30 S201, corresponding to human TFEB S211. HLH-30 peptides identified by MS/MS under basal conditions (CTRL; 20°C) indicated in grey, or after heat shock (HS; 1 h, 36°C) indicated in red. Numbers indicate the amino acid positions of the identified peptides. **(B)** Amino acid sequence of HLH-30 isoform a. Peptides identified by MS/MS are shown in bold, and Ser201 is highlighted in red. **(C)** Fragment ion maps of the S201-containing HLH-30 peptide QVVSSSAPTSSIDIE under CTRL and HS conditions. N-terminal and C-terminal fragment ions are shown below and above the peptide sequence, respectively, and color intensity indicates the number of detected fragment ions. S201 was identified as phosphorylated under CTRL conditions but not following HS. **(D-E)** Representative MS/MS spectra of the Ser201-containing HLH-30 peptide QVVSSSAPTSSIDIE under (D) CTRL and (E) HS conditions. The CTRL spectrum identifies phosphorylation at Ser201, whereas the corresponding peptide detected following HS is unphosphorylated. Detected fragment ions are indicated.

**Supplemental Figure 3.**
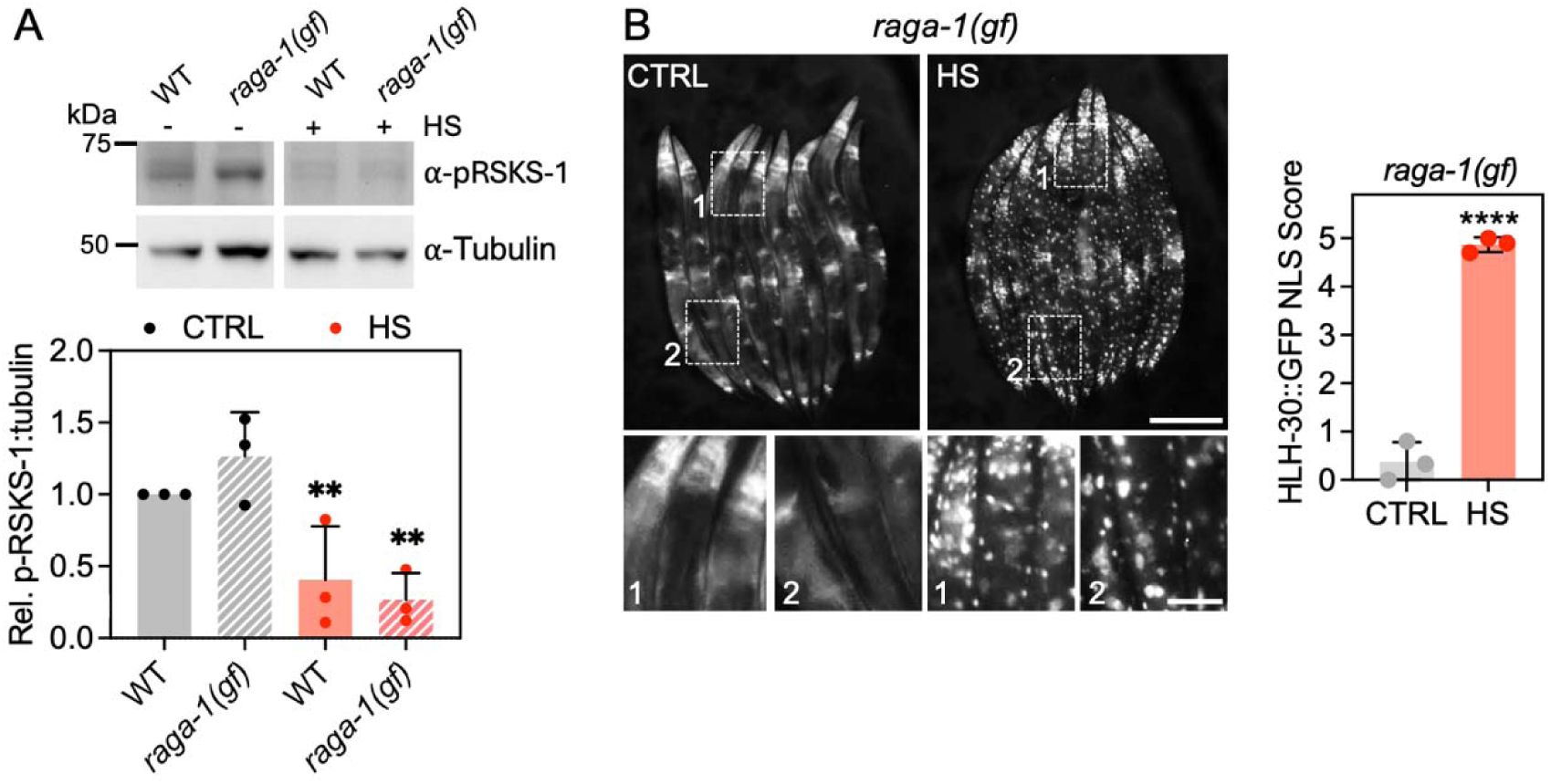
Heat shock inhibits mTORC1 and induces HLH-30 nuclea translocation in *raga-1/RagA* gain-of-function mutants. **(A)** Representative Western blot and quantification of phosphorylated RSKS-1/S6K (pRSKS-1) in wild-type (WT) and *raga-1(gf)* animals under basal conditions (CTRL; 20°C) or following heat shock (HS; 1 h at 36°C). pRSKS-1 levels were normalized to α-tubulin and expressed relative to untreated WT controls. WT CTRL vs. WT HS, P = 0.002; *raga-1(gf)* CTRL vs *raga-1(gf)* HS, P = 0.002; by two-way ANOVA. n = 3. **(B)** Representative images and quantification of HLH-30::GFP subcellular distribution in day 1 adult *raga-1(gf)* mutants under basal (CTRL; 20°C) and heat shock (HS; 36°C) conditions. Scale bar: 200 µm; scale bar in magnified ROIs: 50 µm. **** p<0.0001 by Student’s t-test. n = 3 experiments with N = 10-12 per experiment.

